# Lipopolysaccharide-induced plasma membrane hyperpolarization in *Chara corallina* and *Arabidopsis thaliana*: evolutionary conservation of bacterial elicitor perception across the streptophyte lineage

**DOI:** 10.64898/2026.09.10.750543

**Authors:** Lucie Baudin, Tom Keller de Schleitheim, François Bouteau

**Author notes:** Corresponding author: François Bouteau.

## Abstract

The green alga *Chara corallina* (Charophyceae) is a long-standing model for plant electrophysiology and, owing to its phylogenetic proximity to the ancestors of land plants, an informative system for probing the evolutionary depth of plant–microbe interactions. Bacterial lipopolysaccharide (LPS) is a potent microbe-associated molecular pattern (MAMP) that triggers innate immune signaling in both animals and land plants, but nothing was known about how a charophyte alga responds to this elicitor at the level of the plasma membrane. Here we measured resting membrane potential (V_m_) changes in *C. corallina* internodal cells exposed to purified LPS from three Gram-negative bacteria: *Escherichia coli, Pseudomonas aeruginosa*, and *Pectobacterium carotovorum* subsp. *carotovorum*. All three LPS preparations triggered hyperpolarization of the plasma membrane, but with strikingly different kinetics: *E. coli* LPS produced a transient hyperpolarization that decayed within minutes, whereas LPS from *P. aeruginosa* and *P. carotovorum* produced a sustained, non-transient hyperpolarization that persisted for the duration of recording. Pretreatment with vanadate, a specific inhibitor of P-type H^+^-ATPases, abolished the electrogenic component of the resting potential and strongly reduced the LPS-induced hyperpolarization, suggesting that the plasma-membrane proton pump participates in this response. Strikingly, membrane-potential recordings performed in *Arabidopsis thaliana* cell-suspension cultures reproduced chemotype-specific kinetic dichotomy: LPS from the non-phytopathogen *E. coli* and *P. aeruginosa*, both induced a transient hyperpolarization, whereas LPS from the phytopathogen *P. carotovorum* subsp. *carotovorum* induced a non-transient, sustained hyperpolarization of *A. thaliana* cells. These results demonstrate that *Chara* possesses an electrophysiologically detectable LPS-sensitive perception system that pre-dates the divergence of land plants, that this system can discriminate between LPS chemotypes with distinct kinetic signatures, and that the same transient/non-transient signature is conserved in *A. thaliana*. These findings argue that the transient/non-transient dichotomy documented here may reflect an ancestral, broadly conserved capacity to decode structural variation in bacterial glycolipids at the level of the plasma membrane.

## Introduction

*Chara corallina* and related Characeae have served as a reference system in membrane electrophysiology since the earliest intracellular recordings of resting and action potentials in plant cells (Beilby 2019). The giant, multinucleate internodal cells of *Chara* combine a size that is readily amenable to microelectrode impalement, voltage clamping and vibrating-probe ion-flux measurements, with an ion-transport machinery, voltage-gated Ca^2+^ channels, Ca^2+^-activated Cl^−^ channels, and a vanadate-sensitive plasma-membrane H^+^-ATPase, that is broadly homologous to that of land plants (Kataev et al. 2012; Zhang et al. 2016). Because Characeae belong to the streptophyte algal lineage that gave rise to embryophytes, and because the genome of *Chara braunii* has revealed extensive conservation of genes once thought to be land-plant innovations (Kurtović et al. 2024), *Chara* offers a unique opportunity to ask whether physiological responses considered hallmarks of land-plant immunity already existed, in some form, before the water-to-land transition.

*Chara corallina* cells do not live in isolation from bacteria. Their cell walls and surfaces support dense and taxonomically structured epiphytic bacterial biofilms (Hempel et al. 2008), and at least one Gram-positive pathogen, *Bacillus cereus*, actively attacks the *Chara* plasma membrane through the pore-forming toxin HlyII, producing measurable decreases in resting potential and membrane resistance (Kataev et al. 2012). These observations establish that *Chara* plasma membrane electrophysiology is sensitive to bacterial products, but they concern proteinaceous cytolysins rather than the glycolipid MAMPs that dominate innate-immune signaling in both the animal and plant kingdoms. Lipopolysaccharide (LPS), the outer-membrane glycolipid of Gram-negative bacteria, is one of the most extensively studied MAMPs. In mammals, the lipid A moiety of LPS is sensed by Toll-like receptor 4 and triggers pro-inflammatory signaling. In *A. thaliana*, LPS was originally reported to be perceived by the S-domain lectin receptor-like kinase LORE (LipoOligosaccharide-specific REduced elicitation), and *lore* mutants show increased susceptibility to *Pseudomonas syringae* (Ranf et al. 2015). Subsequent work refined this model considerably: LORE does not bind intact LPS but rather free medium-chain 3-hydroxy fatty acids, which are released as by-products of lipid A biosynthesis and contaminate many LPS preparations (Kutschera et al. 2019). Whether any comparable perception capacity exists outside angiosperms, and in particular within the streptophyte algae, has, to our knowledge, never been addressed electrophysiologically.

In the study of Kettani-Halabi et al. (2015), we directly compared LPS from the phytopathogen *P. carotovorum* subsp. *carotovorum* with LPS from the non-phytopathogens *E. coli* and *Pseudomonas aeruginosa* in *Arabidopsis thaliana* cell suspensions. That study established, at the biochemical and transcriptional level, that the two LPS classes are indeed perceived differentially: only *Pectobacterium* LPS triggered a dose-dependent, metabolically active, actinomycin D/cycloheximide-sensitive programmed cell death, with an early and comparatively modest reactive-oxygen-species (ROS) burst peaking around 1 h post-elicitation; LPS from *E. coli* and *P. aeruginosa*, in contrast, did not trigger cell death and instead produced a delayed but larger ROS burst peaking around 3 h, together with a marked alkalization of the extracellular medium that *Pectobacterium* LPS failed to elicit. Both defense-gene responses (*PAL1, PR1*) were ROS-dependent and NADPH-oxidase-dependent for all LPS types, but with chemotype-specific kinetics, consistent with the idea that *Arabidopsis* cells recognize structural features of LPS, likely located in the lipid A and/or core-oligosaccharide moieties, which are known to differ subtly between *Pectobacterium* and *E. coli*, that are not broadly conserved across Gram-negative bacteria.

We therefore tested the hypothesis that *C. corallina* internodal cells possess a plasma-membrane response to bacterial LPS, and that this response is sensitive to the origin of the LPS. Because we previously reported LPS-chemotype discrimination in *A. thaliana* cell suspensions at the biochemical level (Kettani-Halabi et al. 2015), we extended our membrane-potential recordings to *A. thaliana* cultured cells challenged with the same LPS preparations, to test directly whether this discrimination could be observed through electrophysiological responses.

## Materials and Methods

### Plant material

*Chara corallina* was cultured in artificial pond water (APW: 0.1 mM KCl, 1.0 mM NaCl, 1.0 mM CaCl_2_, 1.0 mM HEPES/NaOH, pH 7.4) under a 14/10h light/dark regime, following conditions comparable to those used in previous *Chara* electrophysiological studies (Kataev et al. 2012; Zhang et al. 2016). Single internodal cells (0.7–0.8 mm diameter, 40–50 mm long) with intact node cells were isolated a few hours before experiments.

*Arabidopsis thaliana* (ecotype Columbia) cell suspension cultures were maintained in Gamborg medium supplemented with sucrose and α-naphthalene acetic acid, under continuous shaking, and sub-cultured weekly, following the conditions used by previous studies (Reboutier et al. 2002; Tran et al. 2013; 2018; Kettani-Halabi et al. 2015). Cells in log phase (4 d after sub-culturing) were used for experiments.

### LPS preparations

LPS from *E. coli* (serotype O111:B4), *Pseudomonas aeruginosa* (serotype 10), and *Pectobacterium carotovorum* subsp. *carotovorum* (Pcc) were prepared as in Kettani-Halabi et al. (2015) and applied to *C. corallina* and *A. thaliana* cultured cells at 100 µg·mL^−1^, the concentration at which biochemical LPS responses were previously characterized in various plant cells. Purified LPS from *E. coli, P. aeruginosa*, and *P. carotovorum* subsp. *carotovorum* were dissolved in distilled water and applied to the external bathing medium.

### Electrophysiology

Resting membrane potential of *C. corallina* internodal cells and individual *A. thaliana* cultured cells was recorded by intracellular impalement with glass microelectrodes (Clark GC 150F; Clark Electromedical, Pangbourne, Reading, UK) pulled on a vertical puller (Narishige PEII). Their tips were < 1µm in diameter; they were filled with 600 mM KCl. Impalement was carried out with a micromanipulator (PCS-5000; Burleigh Inst., Hampshire, UK) in a chamber (1 mL) made of Perspex. Experiments were carried out at room temperature (20 to 30°C in june 2026). Electrodes were connected to an Axoclamp 2B amplifier.

For *C. corallina* impalement were done in SM5 medium (0.1 mM KNO_3_, 0.1 mM MgCl_2_,0.1 mM CaCl_2_, 2.0 mM MES adjusted with NaOH to pH 5.5) after a few hours of adaptation. For *A. thaliana* impalement were done on 4-d-old cells maintained in their culture medium to limit stress (main ions in the medium after 4 d of culture: 9 mM K^+^, 11 mM NO_3_, Reboutier et al., 2002). Individual cells were immobilized by a microfunnel and controlled by a micromanipulator (WR6-1; Narishige, Herts, UK).

## Results

In this experimental series, no *Chara* cell displayed the strongly negative resting potentials (below −150 mV) classically reported for *Chara* internodal cells; all cells were, instead, weakly polarized, with a mean baseline V_m_ around −38 ± 2.7 mV (n=16), consistent with a “leak state” dominated by passive anion conductance. Cells in this state were retained rather than excluded, because *A. thaliana* cell-suspension cultures, the comparator system used in this study, are themselves known to sit, under standard culture conditions, in a comparably weakly polarized, anion-conductance-dominated electrical state, with resting potentials likewise around −40 mV (Reboutier et al. 2002; Tran et al. 2013; 2018). Recording both species under this matched electrical regime, rather than restricting *Chara* recordings to the more strongly polarized state cells reported in other studies, was considered a methodologically consistent basis for the cross-species comparison developed in this study. Moreover, these *Chara* internodal cells maintain a substantial vanadate-sensitive component of their resting conductance (Fig.1), consistent with the well-established role of a P-type H^+^-ATPase in setting the *Chara* resting potential (Beilby 2019; Zhang et al. 2016), since treatment with vanadate (500 µM) allowed decreasing the membrane potential of 13.3 ± 2.3 mV (n=7) mV indicating an electrogenic component dependent of a P-type H^+^-ATPase.

**Figure 1:**
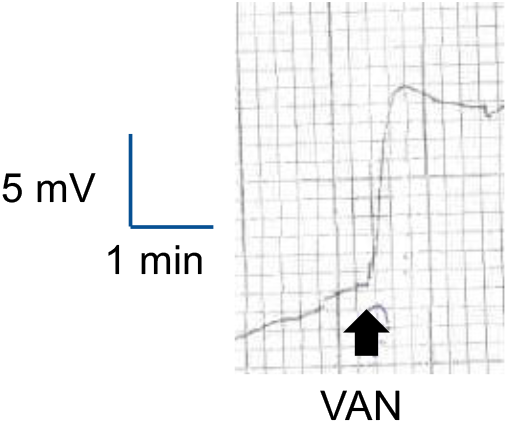
Typical depolarization of Chara cell induced by 500 µM vanadate.

### All three LPS chemotypes hyperpolarize the *C. corallina* plasma membrane

Application of LPS from *E. coli, Pseudomonas* sp. or *Pectobacterium* sp. to the external medium consistently produced a rapid hyperpolarization of the *C. corallina* plasma membrane (30/34 tests), confirming that internodal cells are electrically responsive to Gram-negative bacterial glycolipids as a class. LPS-induced hyperpolarization, including the chemotype-specific transient/non-transient kinetics described below, was recorded from this uniformly weakly polarized cell population (Fig. 2A,C).

**Figure 2:**
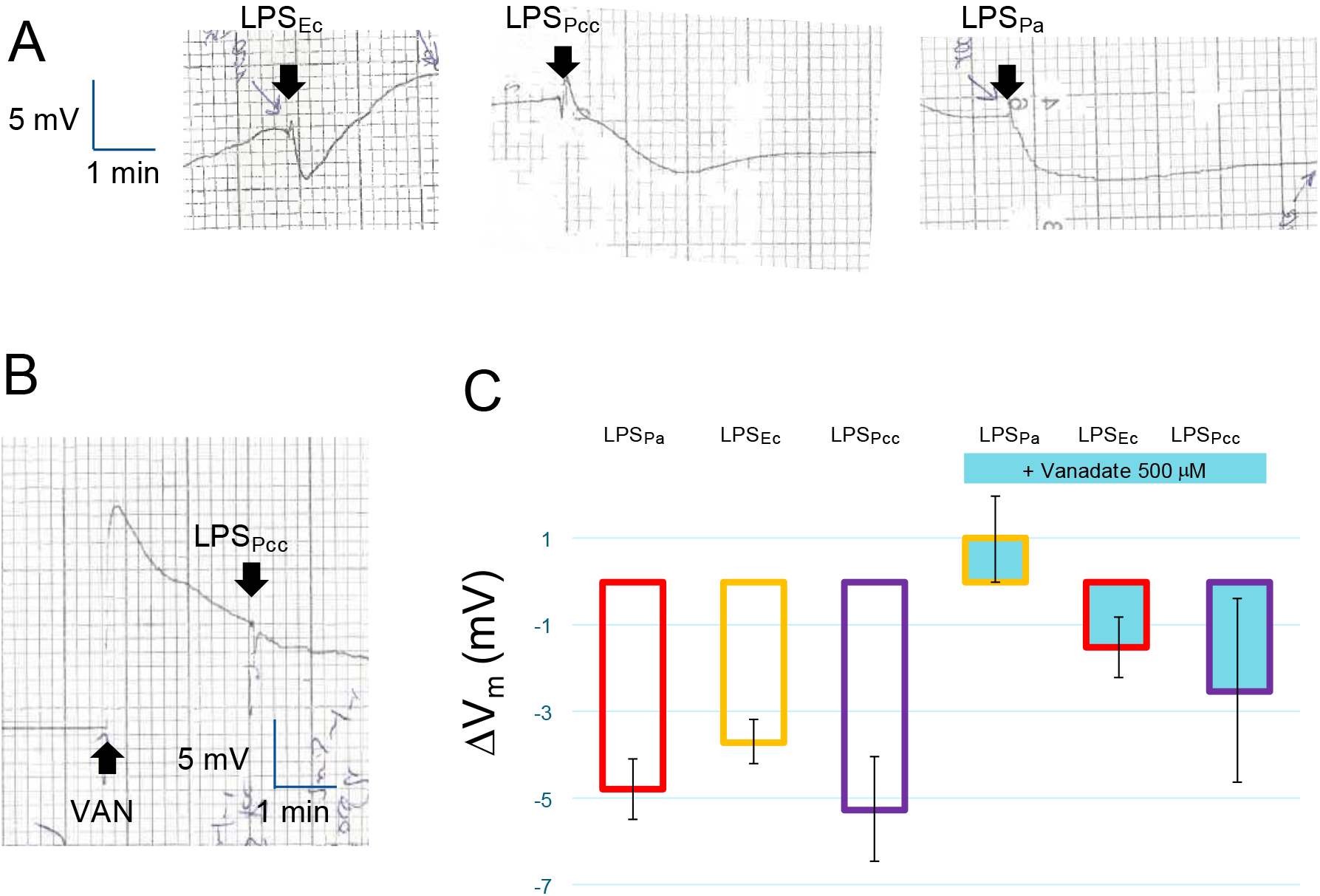
**A**. Hyperpolarizations induced by various LPS on Chara cells. **B**. Example of the absence of hyperpolarization by LPS_Pcc_ after pretreatment by 500 µM vanadate. **C**. Mean values of the hyperpolarizations induced by LPS on C. corallina cells. Blue bars represent membrane potential variation induced by LPS in presence of 500 mM vanadate. Data from at least seven independent cells per condition were averaged for LPS effects and 3 cells for LPS plus vanadate effects.

Following addition of LPS_Ec_, V_m_ hyperpolarized within the first minutes of exposure and then relaxed back towards the pre-stimulus baseline, typically within 1–2 min, despite continued presence of the elicitor in the bath (Fig. 2A). This transient kinetic profile could be reproduced on the same cells suggesting no desensitising. In marked contrast, LPS from *P. aeruginosa* and from *P. carotovorum* ssp. *carotovorum* produced a hyperpolarization of comparable or greater initial amplitude that did not relax back to baseline (Fig. 2A); V_m_ remained persistently hyperpolarized for the entire recording period that could last 10 min. The central result of this study is therefore the clear kinetic dichotomy between the *E. coli* LPS response (transient hyperpolarization) and the *Pectobacterium*/*Pseudomonas* LPS response (non-transient, sustained hyperpolarization), obtained under otherwise identical experimental conditions. However, the extents of the hyperpolarization are of the same order of magnitude for the different LPS (Fig. 2C).

### The hyperpolarizing response depends on a vanadate-sensitive electrogenic pump

Pre-treatment with vanadate abolished or strongly reduced the hyperpolarizing responses to all three LPS preparations, irrespective of whether the untreated response had been transient (*E. coli*) or non-transient (*P. aeruginosa, P. carotovorum*) (Fig. 2B,C). This suggests that the plasma-membrane H^+^-ATPase could be the common downstream electrogenic effector for LPS-triggered hyperpolarization in *Chara*, while the upstream events that determine the transient versus sustained kinetics of activation differ according to LPS origin.

### The transient/non-transient dichotomy is conserved in *A. thaliana* cells

To test whether the kinetic dichotomy observed in *C. corallina* has a counterpart in an angiosperm, we recorded membrane potential in *A. thaliana* cell-suspension cultures challenged with the same panel of LPS. As previously reported (Reboutier et al. 2002; Tran et al., 2013; 2018) in their cultured medium, cultured cells display weak values of resting membrane potential (-40 ± 5 mV, n = 25). However, in these conditions, addition of LPS from *E. coli* induced a hyperpolarization of the *A. thaliana* cells plasma membrane that was transient, relaxing back towards the pre-stimulus baseline within a time-course a little bit longer to that seen in Chara (Fig. 3A). In marked contrast, LPS from the phytopathogen *P. carotovorum subsp. carotovorum* induced a non-transient, sustained hyperpolarization that did not recover within the recording period (Fig. 3A). Unexpectedly, LPS from the non-phytopathogenic bacterium *P. aeruginosa* also produced a transient hyperpolarization in *A. thaliana* cells (Fig. 3A), closely matching the kinetics obtained with *E. coli* LPS and diverging from the sustained profile obtained on *Chara*. These data establish for the first time, a direct electrophysiological read-out of LPS-chemotype discrimination in *A. thaliana* cells: transient hyperpolarization for LPS_Ec_ and LPS_Pa_, and non-transient hyperpolarization for LPS_Pcc_. This chemotype-specific pattern resembles that recorded in *C. corallina* since different types of hyperpolarization are observed with the exception of LPS_Pa_ that induced different type of hyperpolarization according to the models. The extents of hyperpolarization are also in the same range in both models (Fig. 2C,3B).

**Figure 3:**
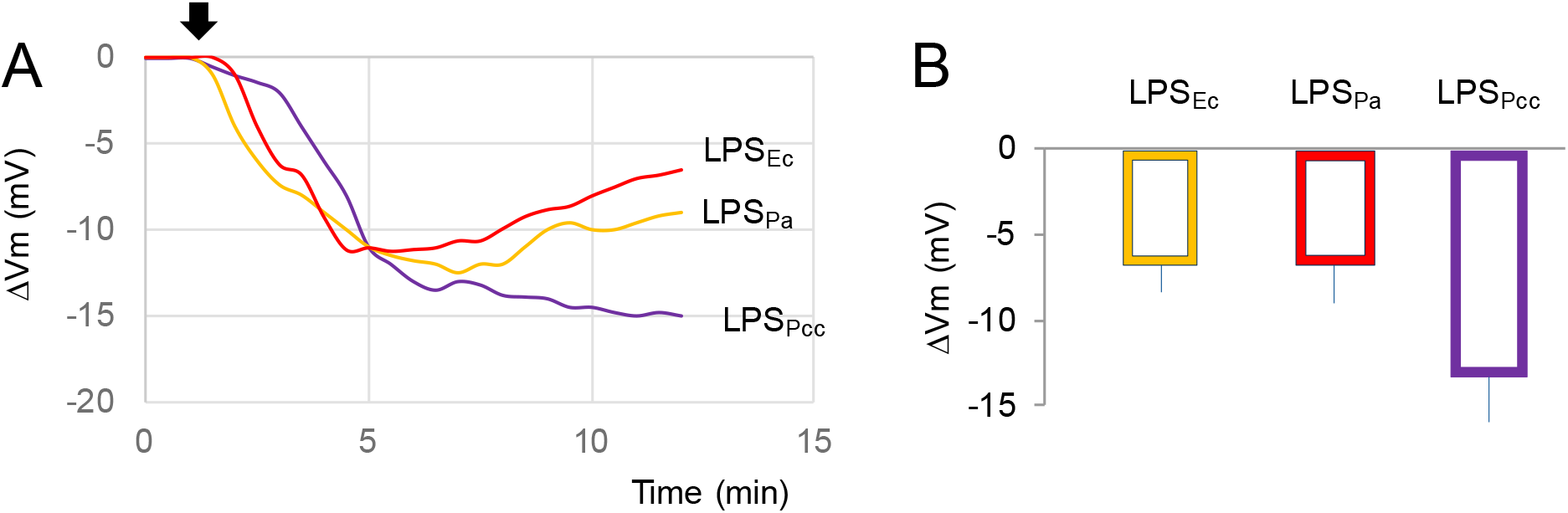
Hyperpolarization induced by various LPS on A. thaliana cultured cells. The arrow indicates the addition of LPS. Mean values of hyperpolarization induced by the various LPS. Data from at least six independent cells per condition were averaged for LPS effects.

## Discussion

### An LPS-sensitive electrical response pre-dates land plants

The observation that *C. corallina*, a streptophyte alga that diverged from the land-plant lineage several hundred million years before the origin of *A. thaliana*, mounts a rapid and reproducible plasma-membrane hyperpolarization in response to bacterial LPS is, to our knowledge, the first electrophysiological demonstration of MAMP-like perception in a charophyte. Because *Characeae* retain many ancestral traits and because the *Chara braunii* genome contains numerous genes previously considered land-plant innovations (Kurtović et al. 2024), this finding is consistent with the view that at least a rudimentary capacity to detect bacterial glycolipids, and to couple this detection to a change in plasma-membrane electrogenesis, was already present in the common ancestor of *Characeae* and land plants.

### The vanadate-sensitive H^+^-ATPase as a conserved electrogenic effector

In *Chara*, the P-type H^+^-ATPase is the principal electrogenic pump responsible for the negative resting potential, and its stimulation is already known to underlie hyperpolarizing responses to other signals, notably auxin (Zhang et al. 2016). Our finding that vanadate reduces LPS-induced hyperpolarization irrespective of LPS origin places this same pump downstream of bacterial elicitor perception. In land plants, elicitor perception classically converges on the plasma membrane H^+^-ATPase as well, although the polarity of the response differs: MAMP perception in angiosperms (e.g., flg22, elf18) typically triggers rapid H^+^-ATPase inhibition and depolarization, whereas we observe H^+^-ATPase-dependent hyperpolarization in *Chara*. This divergence may reflect a fundamental reorganisation of MAMP-triggered signalling polarity during land-plant evolution, or alternatively may indicate that the *Chara* LPS response reported here is mechanistically closer to a growth-promoting or homeostatic electrogenic reflex than to a bona fide defense response. However, MAMPs such harpin from the Gran negative *Erwinia amylovora* were also shown to induce hyperpolarization of the same extent in *A. thaliana* cells (El-Maarouf et al. 2001) suggesting different possible effect of MAMP on H^+^-ATPase regulation, although this response depend on the host/non-host aspect of the plant microbe interaction (Reboutier et al. 2007). This regulation would thus require pharmacological and molecular follow-up.

A methodological point raised by the baseline V_m_ deserves explicit discussion, because it bears directly on the interpretation of the vanadate-sensitive component. Healthy, strongly polarized *Chara* internodal cells are often reported, in other studies, to sit in the “pump state,” with resting potentials well below -150 mV, driven by strong electrogenic activity of the plasma-membrane H^+^-ATPase (Beilby 2019; Zhang et al. 2016). In the present experimental series, however, no cell reached this strongly polarized state: all recorded *C. corallina* cells were, instead, uniformly weakly polarized, with a mean baseline V_m_ around −40 mV. Such a depolarized baseline is itself informative: it is far less negative than would be expected even from a simple loss of pump activity alone (which typically brings V_m_ to an intermediate, diffusion-dominated potential in the −100 to −150 mV range), and instead points to a resting conductance dominated by a non-selective or Cl^−^-dominated leak pathway that clamps V_m_ close to the Cl^−^ equilibrium potential. We chose to retain, rather than discard, this weakly polarized cell population, for a reason directly relevant to the comparative design of this study. *Arabidopsis thaliana* cell-suspension cultures, the system used by Kettani-Halabi et al. (2015) for the biochemical characterization of the same LPS panel, are themselves known to sit, under standard culture conditions, in a comparably weakly polarized, anion-conductance-dominated electrical state, with resting potentials likewise around −40 mV (Reboutier et al. 2002, Tran et al. 2013), well short of the strongly negative, pump-dominated potentials reported in numerous plant cells. Recording *C. corallina* cells in this same weakly polarized, anion-dominated regime, rather than restricting the analysis to the more strongly polarized “pump state” cells sometimes used in other *Chara* electrophysiology studies, therefore places the two species on a matched electrical footing, and removes a potential confound whereby any observed similarity or difference between *C. corallina* and *A. thaliana* could otherwise be attributed simply to the two systems being examined in different, non-comparable baseline states.

This experimental choice does, however, raise a legitimate question: can the LPS-induced, vanadate-sensitive hyperpolarization still be reliably recorded and correctly attributed to the H^+^-ATPase in cells whose baseline conductance is dominated by a passive anion pathway rather than by the pump? Three considerations indicate that it can, while also defining the limits of this interpretation.

First, from simple circuit considerations, the amplitude of any voltage deflection produced by a change in pump current (ΔI_pump) scales with the parallel membrane resistance (ΔV_m_ ≈ ΔI_pump × Rm). A low Rm arising from a dominant anion conductance will therefore attenuate, but not necessarily abolish, a genuine pump-dependent hyperpolarization: provided the LPS-induced increase in pump current is large enough, a measurable hyperpolarization can still be recorded even from a weakly polarized, anion-conductance-dominated cell, simply of smaller absolute amplitude than would be expected from a strongly polarized “pump state” cell. This is fully consistent with our observation that clear, chemotype-specific LPS-induced hyperpolarization was recorded from this uniformly weakly polarized cell population.

Second, and more decisively, two features of the response argue against attributing it to a trivial, LPS-independent relaxation of the anion conductance (e.g., spontaneous closure of Ca^2+^-activated Cl^−^ channels coincidentally following LPS addition) rather than to genuine H^+^-ATPase stimulation. (i) The kinetics of the hyperpolarization are chemotype-specific, transient for LPS_Ec_, sustained for LPS_Pa_/LPS_Pcc_, a pattern that a generic, LPS-independent channel-closing artefact occurring at a fixed intrinsic rate could not reproduce, since it has no means of “knowing” which LPS chemotype was applied. (ii) The hyperpolarization is inhibited by vanadate, a P-type ATPase inhibitor with no established direct action on *Chara* Ca^2+^-activated Cl^−^ channel gating. A channel-closing artefact would not be expected to depend on vanadate. Together, chemotype-specificity and vanadate-sensitivity constitute converging evidence that the hyperpolarization reflects a real, LPS-triggered increase in H^+^-ATPase-mediated electrogenic current, superimposed on the dominant background anion conductance that characterized all cells in this series.

Third, vanadate is not perfectly selective for the plasma-membrane H^+^-ATPase and also inhibits Ca^2+^-ATPases, including those responsible for cytosolic Ca^2+^ clearance at the plasma membrane and tonoplast. In principle, vanadate could therefore blunt the LPS-induced hyperpolarization indirectly, by prolonging a cytosolic Ca^2+^ elevation and thereby sustaining activation of Ca^2+^-activated Cl^−^ channels, rather than by directly blocking an LPS-stimulated H^+^ pump current. This alternative mechanism would still implicate a vanadate-sensitive ATPase in the response, but would revise its precise electrogenic origin. Definitively excluding this possibility, and quantifying the true pump-dependent current independently of the parallel anion conductance, will require voltage-clamp recordings of the vanadate-sensitive instantaneous current-voltage relationship (Kataev et al. 2012), rather than resting V_m_ measurements alone, and ideally a comparison of pre- and post-LPS Rm to determine whether the hyperpolarization is accompanied by an increase (consistent with anion-channel closure) or no change/decrease (consistent with a pump-dominated mechanism) in membrane resistance, in both *C. corallina* and *A. thaliana*.

### Evolutionary comparison with the *A. thaliana* LORE pathway

It is tempting to draw a direct parallel between the LPS-chemotype-specific responses reported here and the *A. thaliana* the lectin receptor-like kinase LORE pathway. However, the comparison must be made carefully. LORE was originally identified as an LPS receptor conferring sensitivity to LPS from *Pseudomonas* and *Xanthomonas* in *A. thaliana*, and *lore* loss-of-function mutants are hypersusceptible to *P. syringae* infection (Ranf et al. 2015). It was subsequently shown that LORE does not directly bind intact LPS but instead recognizes free medium-chain 3-hydroxy fatty acids that are released during lipid A biosynthesis and are present as contaminants in many LPS preparations (Kutschera et al. 2019). The sustained, non-transient hyperpolarization we observe in *C. corallina* upon exposure to LPS_Pa_ and LPS_Pcc_ could, in principle, be driven by the same class of free 3-hydroxy fatty acid contaminants that activate LORE in *A. thaliana*, rather than by the LPS backbone itself, whereas the transient response to LPS_Ec_ may reflect a lower content of such free lipids, or recognition of a distinct structural feature of the *E. coli* lipid A/core-oligosaccharide. Because *Chara* does not seem possessing a LORE orthologue and, more generally, have only o few genes of the receptor-like kinase families that dominate MAMP perception in angiosperms (Dievart et al. 2020), any LPS-sensing capacity in *Chara* must rely on an as-yet-unidentified mechanism, conceivably a lipid-sensing element embedded directly in the plasma membrane or associated with the H^+^-ATPase itself, rather than a dedicated extracellular receptor kinase.

Taken together, an evolutionary scenario in which an ancestral, receptor-independent capacity to translate structural variation in bacterial glycolipids into distinct electrogenic kinetic signatures was present in the common ancestor of *Characeae* and land plants. This capacity appears to have been retained essentially unchanged, at the level of the plasma-membrane electrical signature in *A. thaliana* itself. Under this scenario, the transient/non-transient dichotomy between *E. coli* and *P. carotovorum*/*P. aeruginosa* LPS is not a *Chara*-specific curiosity but the most direct experimental signature currently available of a discriminatory electrogenic capacity conserved across more than 450 million years of streptophyte evolution.

The *Arabidopsis* electrical dichotomy in LPS chemotypes, *E. coli* LPS_Ec_ and LPS_Pa_ produces a transient hyperpolarization, whereas LPS_Pcc_ produces a non-transient, sustained hyperpolarization, is in agreement with conclusions from Kettani et al. (2015) suggested by ROS kinetics, where response to LPS_Pcc_ appeared to be the more transient while LPS_Ec_ and LPS_Pa_ produced a more sustained, delayed ROS response. The membrane-potential data therefore show that the “transient versus sustained” character of LPS action is not a fixed property of the elicitor read out identically by every downstream pathway; rather, the electrical signature and the biochemical (ROS) signature can classify the same LPS chemotypes in opposite ways, presumably because they report on different, only partially coupled branches of the perception cascade.

What is conserved between *C. corallina* and *A. thaliana* is specifically the electrogenic, plasma-membrane-potential branch of the response, not necessarily its downstream biochemical correlates. The result concerning LPS_pa_ was unexpected, which grouped with the non-phytopathogenic, delayed-ROS chemotype in the biochemical study of Kettani-Halabi et al. (2015), but which, in *Arabidopsis* membrane-potential recordings, groups electrically with LPS_Pcc_ as a non-transient hyperpolarizing agent, matching exactly its behaviour in *Chara*. This reinforces the conclusion that pathogenicity status (phytopathogen versus non-phytopathogen) is not, by itself, what the plasma-membrane electrical response is tracking; instead, the electrical read-out appears to track a structural property of the LPS itself, plausibly shared between *Pseudomonas* and *Pectobacterium* lipid A/core-oligosaccharide chemistry, that is independent of the bacterium’s virulence lifestyle on plants, and that is discriminated identically by the electrogenic machinery of a charophyte alga and of an angiosperm.

## Conclusion

Taken together, these results position *C. corallina* and *A. thaliana* not as a divergent pair with respect to LPS-triggered electrogenesis, but as two exemplars of a single, evolutionarily conserved discriminatory mechanism operating at the plasma membrane, detectable by microelectrode-based electrophysiology in both a streptophyte alga and an angiosperm. This substantially strengthens the case that the transient/non-transient electrogenic dichotomy is an ancient feature of the streptophyte lineage rather than a *Chara*-specific idiosyncrasy or an angiosperm-specific innovation. These results suggest that a capacity to perceive and kinetically decode bacterial glycolipids at the level of the plasma membrane evolved before the streptophyte algae diverged from the land-plant lineage and has been retained. Future work should test whether these responses are driven by the LPS backbone itself or by free 3-hydroxy fatty acid contaminants, and should seek to identify the molecular sensor upstream of the H^+^-ATPase in both species. Finally, from an ecological point of view, *Chara* surfaces are naturally colonized by structured bacterial communities that differ between freshwater and brackish habitats and between plant species (Hempel et al. 2008), and *Chara* is also a documented target of Gram-positive pathogenic attack via pore-forming toxins (Kataev et al. 2012). The capacity to electrically discriminate between LPS chemotypes could plausibly allow *Chara* to distinguish commensal or epiphytic Gram-negative bacteria from more aggressive, pectinolytic phytopathogens such as *Pectobacterium*, even in the absence of a dedicated innate-immune receptor repertoire.

## Acknowledgements

*Chara corallina* were kindly provided by Etienne Couturier.

## References

Beilby MJ (2019) Chara braunii genome: a new resource for plant electrophysiology. Biophys Rev 11:235–239.

Dievart A, Gottin C, Périn C, Ranwez V, Chantret N. (2020) Origin and Diversity of Plant Receptor-Like Kinases. Annu Rev Plant Biol. 71: 131–156.

El-Maarouf H, Barny MA, Rona JP, Bouteau F. 2001. Harpin, a hypersensitive response elicitor from Erwinia amylovora, regulates ion channel activities in Arabidopsis thaliana suspension cells. FEBS Lett. 497(2–3):82–4.

Hempel M, Blume M, Blindow I, Gross EM (2008) Epiphytic bacterial community composition on two common submerged macrophytes in brackish water and freshwater. BMC Microbiology 8:58.

Kataev AA, Andreeva-Kovalevskaya ZI, Solonin AS, Ternovsky VI (2012) Bacillus cereus can attack the cell membranes of the alga Chara corallina by means of HlyII. Biochim Biophys Acta 1818:1235–1241.

Kettani-Halabi M, Tran D, Dauphin A, El-Maarouf-Bouteau H, Errakhi R, Arbelet-Bonnin D, Biligui B, Val F, Ennaji MM, Bouteau F (2015) Deciphering the dual effect of lipopolysaccharides from plant pathogenic Pectobacterium. Plant Signal Behav 10:e1000160.

Kurtović K, Schmidt V, Nehasilová M, Vosolsobě S, Petrášek J (2024) Rediscovering Chara as a model organism for molecular and evo-devo studies. Protoplasma 261:183–196.

Kutschera A, Dawid C, Gisch N, Schmid C, Raasch L, Gerster T, Schäffer M, Smakowska-Luzan E, Belkhadir Y, Vlot AC, Chandler CE, Schellenberger R, Schwudke D, Ernst RK, Dorey S, Hückelhoven R, Ranf S (2019) Bacterial medium-chain 3-hydroxy fatty acid metabolites trigger immunity in Arabidopsis thaliana. Science 364:178–181.

Ranf S, Gisch N, Schäffer M, Illig T, Westphal L, Knirel YA, Sánchez-Carballo PM, Zähringer U, Hückelhoven R, Lee J, Scheel D (2015) A lectin S-domain receptor kinase mediates lipopolysaccharide sensing in Arabidopsis thaliana. Nat Immunol 16:426–433.

Reboutier D, Bianchi M, Brault M, Roux C, Dauphin A, Rona JP, Legué V, Lapeyrie F, Bouteau F. (2002) The indolic compound hypaphorine produced by ectomycorrhizal fungus interferes with auxin action and evokes early responses in nonhost Arabidopsis thaliana. Mol Plant Microbe Interact. 15:932–8. doi: 10.1094/MPMI.2002.15.9.932.

Reboutier D, Frankart C, Briand J, Biligui B, Laroche S, Rona JP, Barny MA, Bouteau F. 2007. The HrpN(ea) harpin from Erwinia amylovora triggers differential responses on the nonhost Arabidopsis thaliana cells and on the host apple cells. Mol Plant Microbe Interact. 20(1):94–100.

Tran D, El-Maarouf-Bouteau H, Rossi M, Biligui B, Briand J, Kawano T, Mancuso S, Bouteau F. (2013) Post-transcriptional regulation of GORK channels by superoxide anion contributes to increases in outward-rectifying K+ currents. New Phytol. 198:1039–1048. doi:10.1111/nph.12226.

Tran D, Dauphin A, Meimoun P, Kadono T, Nguyen HTH, Arbelet-Bonnin D, Zhao T, Errakhi R, Lehner A, Kawano T, Bouteau F. (2018) Methanol induces cytosolic calcium variations, membrane depolarization and ethylene production in arabidopsis and tobacco. Ann Bot. 122:849–860. doi: 10.1093/aob/mcy038.

Zhang S, de Boer AH, van Duijn B (2016) Auxin effects on ion transport in Chara corallina. J Plant Physiol 193:37–44.

